# Microclimate Shapes the Phylosymbiosis of Rodent Gut Microbiota in Jordan’s Great Rift Valley

**DOI:** 10.1101/2023.08.01.551195

**Authors:** Enas Al-khlifeh, Sanaz Khadem, Bela Hausmann, David Berry

## Abstract

Host phylogeny and the environment play vital roles in shaping animal microbiomes. However, the effects of these variables on the diversity and richness of the gut microbiome in different bioclimatic zones remain underexplored. In this study, we investigated the effects of host phylogeny and bioclimatic zone on the diversity and composition of the gut microbiota of two heterospecific rodent species, the spiny mouse *Acomys cahirinus* and the house mouse *Mus musculus*, in three bioclimatic zones of the African Great Rift Valley (GRV). We confirmed host phylogeny using the *D-loop* sequencing method and analyzed the influence of host phylogeny and bioclimatic zone parameters on the rodent gut microbiome using high-throughput amplicon sequencing of 16S rRNA gene fragments. Phylogenetic analysis supported the morphological identification of the rodents and revealed a marked genetic difference between the two heterospecific species. We found that bioclimatic zone had a significant effect on the gut microbiota composition while host phylogeny did not. Microbial alpha diversity of heterospecific hosts was highest in the Mediterranean forest bioclimatic zone, followed by the Irano–Turranian shrubland, and was lowest in the Sudanian savanna tropical zone. The beta diversity of the two rodent species showed significant differences across the Mediterranean, Irano–Turranian, and Sudanian regions. The phyla *Firmicutes* and *Bacteroidetes* were highly abundant, and *Deferribacterota, Cyanobacteria* and *Proteobacteria* were also prominent. Amplicon sequence variants (ASVs) were identified that were unique to the Sudanian bioclimatic zone. The core microbiota families recovered in this study were consistent among heterospecific hosts. However, diversity decreased in conspecific host populations found at lower altitudes in Sudanian bioclimatic zone. The composition of the gut microbiota is linked to the adaptation of the host to its environment, and this study underscores the importance of incorporating climatic factors such as elevation and ambient temperature, in empirical microbiome research and is the first to describe the rodent gut microbiome from the GRV.

## Introduction

Geological and climatic variables can have a considerable effect on bioclimatic regions. Species living in the same bioclimatic zone have common evolutionary elements. However, the primary bioclimatic zones on Earth differ to varied degrees in terms of the biotic and abiotic components of the land that support species with similar lifestyles and adaptations. Therefore, bioclimatic zones can provide information on the patterns of hosts and symbionts that coexist with them. The ubiquitous microbial populations in an animal gut, or gut microbiome, is a major partner in most animal symbioses. Therefore, it is important to focus on increasing knowledge of its effects on health and metabolism, and how its hosts respond to stress. Microbiome composition has been found to be irregular, unstable, and not always completely inherited, primarily because of the interaction between modifications in the microbiome ecosystem and changes in the evolutionary past of the host (Brooks et al., 2016). Therefore, the term “phylosymbiosis” has recently gained popularity.

According to the traditional definition of phylosymbiosis, hosts belonging to the same species are more likely to have similar microbiota than those belonging to distinct species. Strongly settled phylosymbiotic patterns are caused by microbial colonization preferences for specific host genetic identities (Lim and Bordenstein, 2020). A portion of microbiome traits have a strong genetic component, such as vertical inheritance (Ferretti et al., 2018) and physical contact between members of one species (Dill-McFarland et al., 2019). However, these mechanisms alone cannot explain the distinctive microbiome makeup or structure of animals. Given that a microbiome can be acquired through environmental acquisition over the course of an animal’s lifespan (Mukherjee et al., 2021), geography and past climatic conditions need to be taken into account. Microbiomes that are abundant in the host habitat and can colonize host niches are more likely to persist from one host generation to the next. Therefore, the associated microbiomes of various animal species living in the same bioclimatic region are expected to be comparable and analogously adapted to bioclimatic zone conditions.

Several studies have led to an understanding of how the environmental distribution affects phylosymbiosis. However, multiple environmental comparisons have shed light on how many biotic and abiotic factors shape selective pressures, some of which are known to promote adaptation of the host microbiome in some ecosystems more than others. For instance, a number of studies on different host linage have provided evidence of geography playing an important role in the formation of the host-associated microbiome without host-specific input (Moraitou et al., 2022; Goertz et al., 2019) (Joakim et al., 2023; Sun et al., 2020). In the case of microbiome evolution across various mouse lineages, Teng et al. (2022) highlighted the opposing roles of host genetics and the environment. They showed that the environment has a greater effect than host species identity (Teng et al., 2022). In addition, the gut microbiome diversity of many rodent species is thought to be influenced by host genetic and geographic differences (Wang et al., 2022). Geographic trends in microbiota composition in human and mouse gut microbiomes have been recognized (Linnenbrink et al., 2013; Suzuki et al., 2019; Rehman et al., 2016). Suzuki et al. (2018) and Goertz et al. (2019) demonstrated that location and latitudinal zonation, including at small spatial scales, significantly influenced the composition of the gut microbiome in wild-type mice (Suzuki et al., 2019; Goertz et al., 2019). These studies have shed light on the impact of abiotic variables on microbiome composition and abundance despite focusing on certain host species. However, studies that compare heterospecific hosts can be beneficial and help to deepen our understanding of the causes of phylosymbiosis.

The Earth is divided into seven main biogeographical zones by climate parameters, such as temperature, concurrent changes in precipitation levels, vegetation, and soil types (Box, 2016). We focused on three zones of the Great Rift Valley (GRV). The GRV landform separates the southern highlands of Jordan into geographical segments that adjoin the Sudanian, Iran–Turanian, and Mediterranean bioclimatic zones (Ababsa, 2014). Although these regions are believed to have the same evolutionary history and geological characteristics, the Sudanian region faces the African continent from the south and has a tropical climate. Meanwhile, the Mediterranean area is located on a slope confronting the northern side of the European continent. Temperate climatic traits are also observed in this zone. Statistics on temperature and solar radiation have shown that the Sudanese biogeographic region receives eight times more solar radiation than the Mediterranean region (Alrwashdeh et al., 2018; Etier et al., 2010), and its mean annual temperature is 10 °C higher (Ababsa, 2014).

Given the unusual existence of different bioclimatic regions in the GRV, the wildlife fauna is exceptionally diverse, with wild rodents comprising a substantial proportion (Amr et al., 2018a). Rodents encounter various conditions in their natural habitats that affect their level of fitness. Rodents are extremely sensitive to changes in temperature and precipitation, particularly in biogeographic areas (RamírezCBautista et al., 2020). Temperature fluctuations in the environment can influence gut microbiota (reviewed in (Sepulveda and Moeller, 2020a)). Therefore, it is anticipated that climatic variables are likely to have a significant effect on the microbiomes of these animals. Animals demonstrate their own mechanisms of adaptation when confronted with difficult biotic or abiotic circumstances. However, they also depend on their associated symbionts for survival. Moeller et al. (2019) offered experimental evidence that the gut microbiome fueled the adaptive evolution of the house mouse *M. m. domesticus* and had a significant effect on host fitness (Moeller et al., 2019). Rodents are an appropriate model for studying phylosymbiotic patterns in relation to bioclimatic sites.

The spiny mouse *Acomys cahirinus* and the house mouse *M. m. domesticus* are two common rodent species, with a broad distribution range across the three biogeographical regions found in the GRV. Both species has long been used as a genetic model for human biology and disease (Wang et al., 2020) (Gawriluk et al., 2020). *Mus* species have been developed into a model system for microbiota studies, fostering studies on many aspects from the environmental influence on the microbiome to the evolutionary adaptation of microbiota (e.g., (Moeller et al., 2019; Rosshart et al., 2017)). *A. cahirinus* is regarded as a superb model for tissue regeneration and has shed light on the physiological changes brought on by environmental variables (Gawriluk et al., 2020; Haughton et al., 2016). Both species are widely distributed from Western Asia to Africa, including the Mediterranean region, which is thought to be a site where rodents first evolved (Dalecky et al., 2015; Aghová et al., 2019). *Acomys* species, have developed independently from *Mus* for at least 8 Ma (Lecompte et al., 2008). However, both species have been used in several comparative studies on fibrosis and regeneration (Gawriluk et al., 2020; Okamura et al., 2021). Given the phylogenetic distance, it is particularly challenging to compare differentially expressed transcriptomes between *Acomys* and *Mus* for any given phenotype (Okamura et al., 2021; Brant et al., 2019). Therefore, we assumed that these would be excellent test models for determining how the environment affects phylosymbiosis in natural habitats.

In this study, we examined the roles of bioclimatic zone and phylogenetic relatedness (hereon “phylogeny”) as a driver of microbiome structuring in *A. cahirinus* and *Mus. domesticus*. We specifically sought to investigate whether (1) the gut microbiome’s diversity and richness are associated with bioclimatic regions; (2) the host phylogeny can predict the composition of the gut microbiome; and (3) the change in the gut microbiome has fitness implications for its hosts. Here, we considered the body weight of the individuals collected as a proxy for fitness because it is a key component of mouse fitness in the wild (Schulte-Hostedde et al., 2001).

Knowing that the GRV has recorded climate change events that have caused plants to change from the C3 to C4 type (Uno et al., 2016), we proposed that the shift in ambient temperature from the Mediterranean side of the GRV toward a significant elevation in the Sudanian zone represented a reasonable simulation of how climate change may affect the animal gut microbiota. This, in turn, may help us predict how animal lineages and communities respond to future climate change.

## Materials and Methods

### Study site

Owing to the opposing tropical and temperate zones that cross the GRV end in southern Jordan, the study region included biotic and abiotic opposites. Figure 1 depicts the erosion landforms of the valley that crosses the Sudanese and Mediterranean bioclimatic zones. A semi-arid rainforest with a temperate, colder, and mesic microclimate can be found in the Mediterranean zone at greater heights at 700–1500 m above sea level, facing the European continent from the north. In comparison, the Sudanian zone, which faces Africa from the south, has subtropical *Acacia* vegetation and an annual precipitation of less than 50 mm. The valley’s mid altitude steppe flora, which can be found at elevations between 500 and 700 m above sea level, provides evidence of an Irano-Turanian environment. The environmental conditions and soil of each zone are listed in Table 1. Climate analysis has shown that the amount of solar radiation received on the Sudanese side is up to eight times greater than that in the Mediterranean region, and the mean annual temperature is ten degrees higher. These elements help create tropical weather in the area, which is only a few kilometers away from the Mediterranean site.

**Figure 1.**
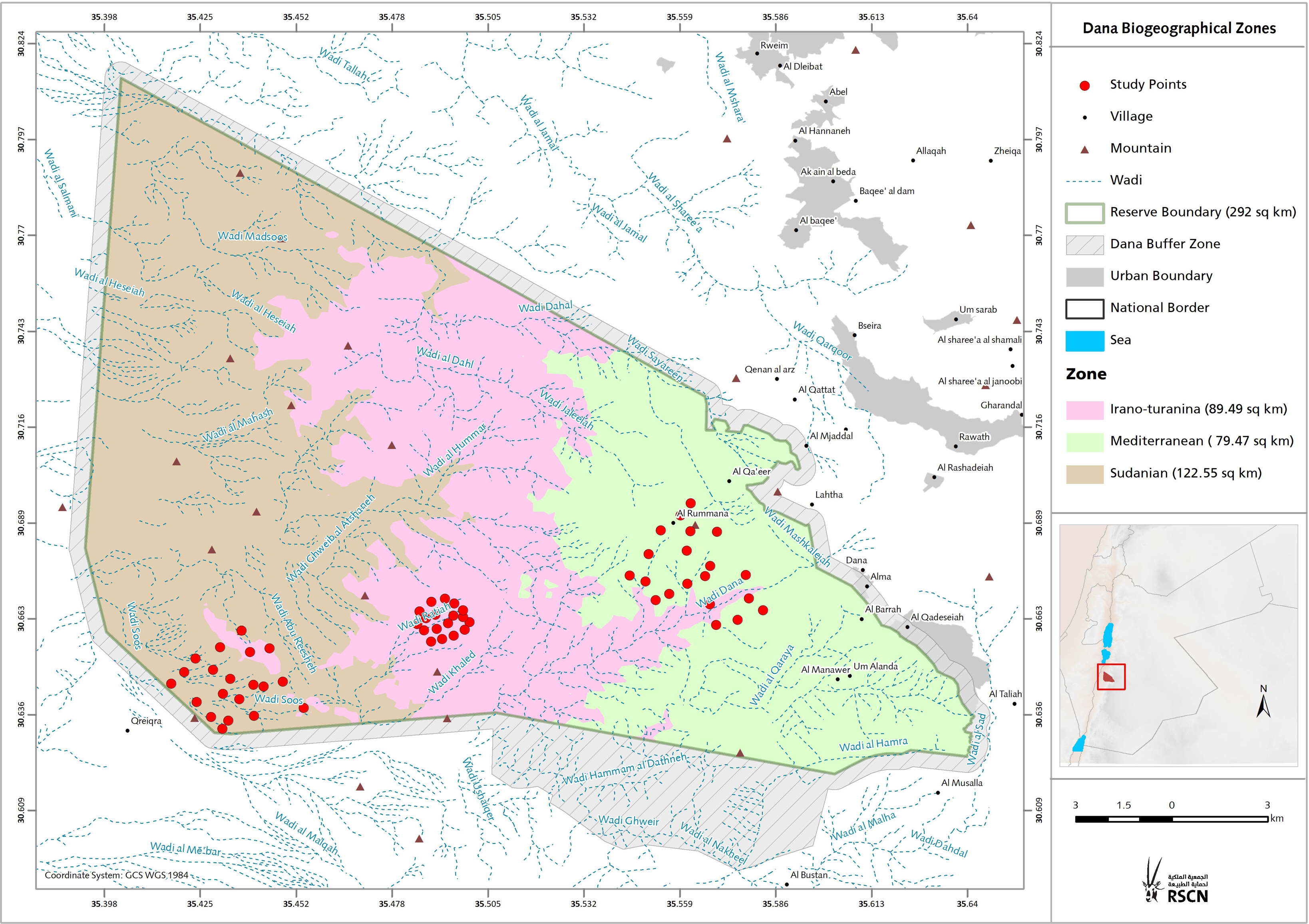
Study Area. Topographic map of the study area showing bioclimatic zones with altitudinal range. The location of the investigation area illustrates the sampling spots in the three bioclimatic zones covers the north-western part of the Arabian plate separated from the African Plate by the Jordan - African Rift Valley.

**Table 1.**
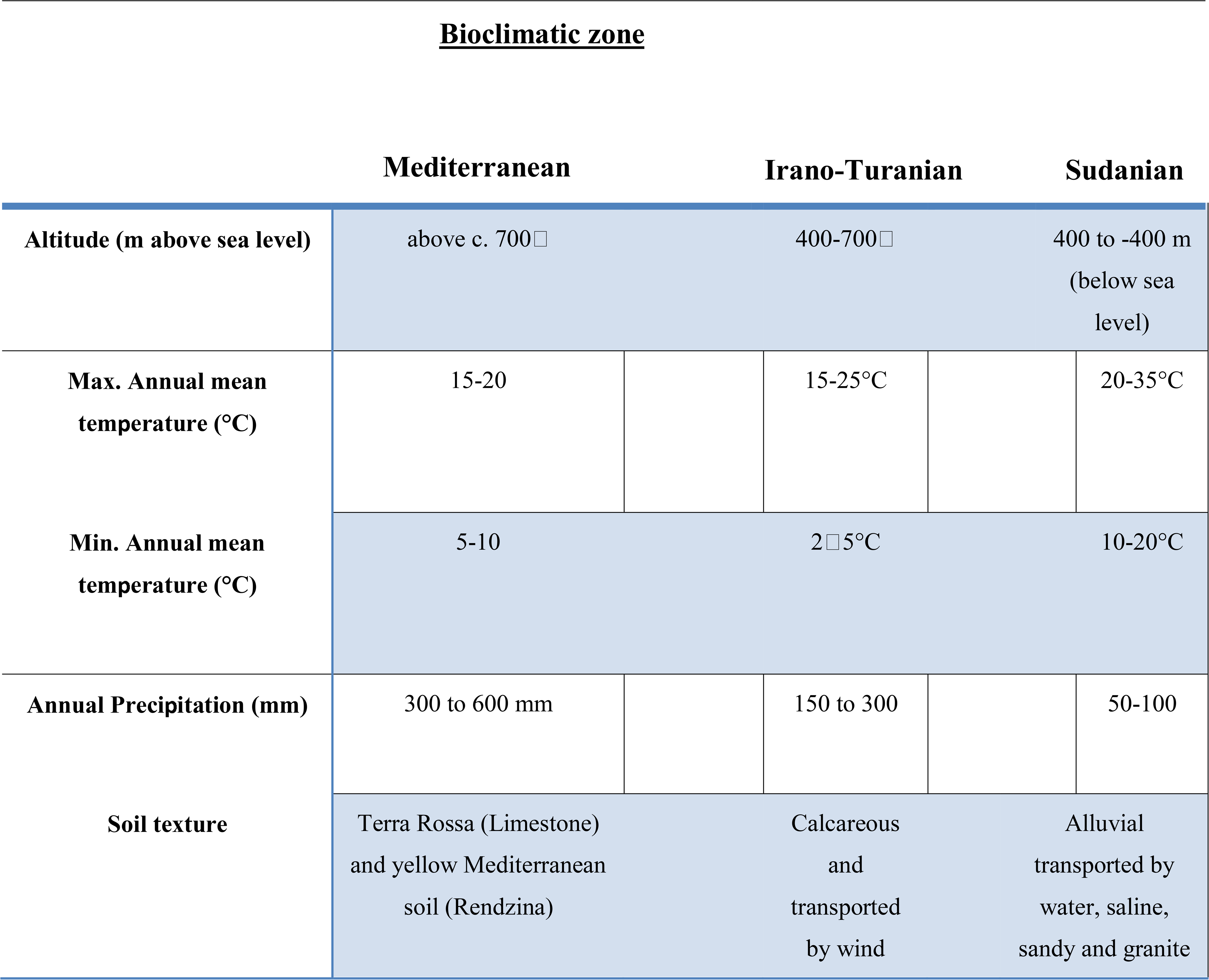
Bioclimatic zone, mean annual temperature, precipitation, and soil at the study sites.

### Rodent trapping and sampling

*Mus mus. domesticus* and *A. cahirinus* and are not on the roster of conserved species because of their abundance. To minimize the dietary changes commonly associated with seasonal changes in the microbiome, sampling was conducted between autumn 2020 and spring 2021. We used Sherman traps positioned 10–20 m apart. Collection sites were chosen randomly because of the geographical barriers. The traps were strategically positioned around rocky holes and crakes, keeping an eye out for active burrows. Traps containing peanut butter and jam were set up at night and tested every morning for 1–3 d at each location before being moved to a new location within the same sampling range. Prior to DNA barcoding, species identification of animals during trapping relied on morphology, as previously described (Amr et al., 2018b). At the time of collection, the weight and reproductive status of each animal were recorded. Any female with signs of pregnancy were excluded. The rodents were euthanized by cervical dislocation immediately after anesthesia. Organs, including the cecum pouch, were stored in Zymo’s DNA/RNA Shield, that is, a preservative medium for biological samples, until DNA isolation. The rodents were captured and handled with the approval of the Jordanian Royal Society for the Conservation of Nature (RSCN).

### Rodent genotyping and phylogeny

Following the manufacturer’s instructions, genomic DNA was recovered from tissue samples using the Invitrogen PureLink Genomic DNA Mini Kit. The nucleotide sequences of the primers used are La1 5′-ATAAAAATTACTCTGGTCTTGTAAAC-3′ (NICOLAS et al., 2009); Bis2 5′-CACAGTTATGGAAGTCTTGG-3′ from (Bellinvia, 2004). The PCR mixture had a total volume of 50.0 µL, which included 2.0 µL of template DNA (10 ng), 5.0 µL of primers, 18.0 µL of deionized distilled water, and 25.0 µL of 2X PCR master mix (i-MAX II) prepared from iNtRONs biotechnology, Inc. Negative controls were used in parallel with PCR amplifications. A thermocycler (BIOER XP Cucler) was used to perform the amplification of a 569 bp *D-loop* fragment. These PCR conditions were used: 15 min at 95 °C and then 35 cycles with 30 s at 95 °C, 1:30 min at 54 °C, and 1 min at 72 °C, with the final elongation lasting 15 min at 72 °C. The genomic DNA concentration was measured using a NanoDrop (Nanodrop Technologies, Thermo Scientific, Waltham, MA, USA). Following the manufacturer’s instructions, Zymo Research DNA Clean & Concentrator® reagents were used to purify the PCR amplicons and remove excess primers and nucleotides. PCR products of appropriate quality and purity were sequenced with Sanger Sequencing (Microsynth, Switzerland). The sequences obtained were assembled and edited manually using the BioEdit software. The Nucleotide Basic Local Alignment Search Tool (blast.ncbi.nlm.nih.gov) of the National Center for Biotechnology was used to search for the acquired consensus sequences to identify their genotype/strain.

MEGA software version 11.0.13 was used for alignment (Tamura et al., 2021), pairwise distance calculation, and drawing the main phylogenetic trees based on the neighbor-joining technique (Saitou and Nei, 1987). We created maximum parsimony trees for both rodent species to gain more understanding of the two separated species (Yang, 1994). To calculate the phylogenetic relationships based on maximum likelihood (ML) method (number of bootstraps = 1000), the PhyML(v3.0) online source was used (Guindon et al., 2010), and iTOL v6.7.6 (https://itol.embl.de/) was performed to visualize the findings. For phylogeny inference, the nucleotide sequences of the isolates examined in this research were aligned with GenBank-retrieved sequences based on complete *D-loop* mitochondrial genomes and chosen outgroup: *Acomys cahirinus* NC020758.1, *Deomys ferrugineus* FJ415539.1, *Acomys dimidiatus* FJ415545.1, *Acomys subspinosus* FJ415548.1, *Acomys spinosissimus* FJ415547.1, *Acomys russatus* MH044885, *Acomys wilsoni* MH044874.1, *Apodemus mystacinus* AY623063, *Mus musculus domesticus* NC 006914.1, *Mus musculus musculus* NC 010339.1, *Mus musculus castaneus* NC 012387.1.

### *16S* rRNA gene sequencing

Genomic DNA extracted from the cecal content material is performed using the QIAamp® Fast DNA Stool Mini Kit according to the manufacturer’s instructions. Target gene amplification and sequencing were performed at the Joint Microbiome Facility of the Medical University of Vienna and the University of Vienna (project IDs JMF-2106-01 and JMF-2209-12) according to the procedure described by Pjevac et al. (2021) (Pjevac et al., 2021). Shortly, 16S rRNA gene amplicons were generated using primers targeting the V4 hypervariable region of most bacteria and archaea (515F/806R; (Apprill et al., 2015; Parada et al., 2016) modified to contain a 16 nt linker overhang each. After the PCR, amplicons were purified and normalized with the SequalPrep Normalization Plate Kit (Invitrogen) following the manufacturers’ instructions, and used as template for a second 8-cycle PCR step, in which each amplicon was barcoded with two unique 12 nt barcode sequences (i.e. unique dual barcoding). Barcoded amplicons were again purified and normalized with the SequalPrep Normalization Plate Kit (Invitrogen), pooled, and concentrated with the innuPREP PCRpure Kit (Analytik Jena). From the amplicon pools, sequencing libraries were prepared with the TruSeq Nano DNA Library Prep Kit (Illumina), excluding the DNA fragmentation step, and sequenced on an Illumina MiSeq using the MiSeq Reagent kit v3 (Illumina, 600 cycles). Individual amplicon pools were extracted from the raw sequencing data using the FASTQ workflow in BaseSpace (Illumina) with default parameters. Demultiplexing was performed with the python package demultiplex (Laros JFJ, github.com/jfjlaros/demultiplex) allowing one mismatch for barcodes and two mismatches for linkers and primers. ASVs were inferred using the DADA2 R package version 1.14.1 Callahan et al. (2016a) with R version 3.6.1 (R Core Team, 2021) applying the recommended workflow (Callahan et al., 2017). Therefore, amplicon FASTQ reads one and two were trimmed at 220 nt and 150 nt with allowed expected errors of 2 and 2, respectively. ASV sequences were subsequently classified using SINA version 1.6.1/1.7.2 (Pruesse et al. 2012) and the SILVA database SSU Ref NR 99 release 138.1 (Quast et al. 2012) using default parameters.

### *18S* rRNA gene sequencing

Amplicon sequencing and raw data processing are carried out utilizing the previously reported two-step PCR barcoding method (Pjevac et al., 2021). The primers used to amplify the hypervariable region of the 18S gene were Next.For (5′ CCCAGCASCYGCGGTAATTCC C 3′) and Next.Rev (5′ CACTTTCGTTCTTGATYRATGA C 3′) (Piredda et al., 2017); (Stoeck et al., 2010). Both primers contained a16nt head sequence at the 5’ to allow further multiplexing, as described in Pjevac et al. (2021). The normalized library was prepared by adapter ligation and PCR using the TruSeq Nano DNA Library Prep Kit according manufacturer’s instructions and subsequently sequenced on a Illumina Miseq platform, v3 2x 300bp (Illumina). Demultiplexing was performed with the python package demultiplex (Laros JFJ, github.com/jfjlaros/demultiplex) allowing one mismatch for barcodes and two mismatches for linkers and primers. Amplicon sequence variants (ASVs) were inferred using the DADA2 R package v1.26 (Callahan et al., 2016a) applying the recommended workflow (Callahan et al., 2016b). FASTQ reads 1 and 2 were trimmed at 240 nt and with allowed expected errors of 4 and 6 respectively. ASV sequences were subsequently classified using DADA2 against the SILVA 18S database with default parameters (Morien and Parfrey, 2018).

### Statistical analysis

All statistical analyses were done in R studio software (https://www.r-project.org/, v4.2.1) and graphs created using ggplot2 package (v3.4.2) (Wickham, 2016). Differences in the number of reads among cecum samples were accounted for by dividing each sequence count by the total number of reads in that sample, yielding relative abundance measures, except in the case of differential expression analysis, for which raw sequence counts were used. 16S libraries were rarefied to a read depth of 5000 reads and using rrarefy function in vegan package (v2.6.2) (Oksanen et al., 2015), eleven samples were excluded from the study because of inadequate reads. Alpha diversity was calculated as observed ASV richness, Shannon diversity index, and Inverse Simpson diversity index. To illustrate beta diversity of microbial compositions, function metaMDS, method “bray” in vegan package (v2.6.2) was used. ASV accumulation curves were obtained with the specaccum function in vegan package (v2.6.2) and indicator ASVs for zones was determined using randomForest (v.4.7.1.1) (Liaw and Wiener, 2002) and visualized with ComplexHeatmap (2.15.1) packages (Gu, 2022), respectively. Enrichments of taxa at bioclimactic zones was calculated with ALDEx2 (ALDEx2 package, v1.30.0) (Gloor et al., 2019).

## Results

### Host phylogeny analysis

At the study site, 120 rodents were captured between October 2020 and May 2021 from each of the three bioclimatic zones. Fifty animals were from the Mediterranean region, 38 from Sudan, and 32 from Iran and Turania. To confirm the taxonomic status and evaluate the population structure based on maternal lineage, we sequenced 569 bp of the mitochondrial *D-loop* region of all 120 mice. Nine of the 120 samples were excluded from the phylogenetic analysis because they were too short or provided ambiguous sequence reads, and two of the 120 samples failed the sequencing process. The *D-loop* ML tree supported the morphological identity of the rodents and revealed a marked genetic difference between samples from *Mus musculus domesticus* and *Acomys cahirinus* (Figure 2). 75 samples were grouped together and considered a single clade of *A. cahirinus*. This group was referred to as the *Acomys* clade. The remaining 34 specimens formed distinct clades within the *M. m. domesticus* superclade. Although both species were detected in all three bioclimatic zones, there was an observable difference in their distribution (Chi-square test, *p*<0.01), with *M. m. domesticus* most prevalent in the Sudanian and *A. cahirinus* in the Mediterranean bioclimatic zone.

**Figure 2.**
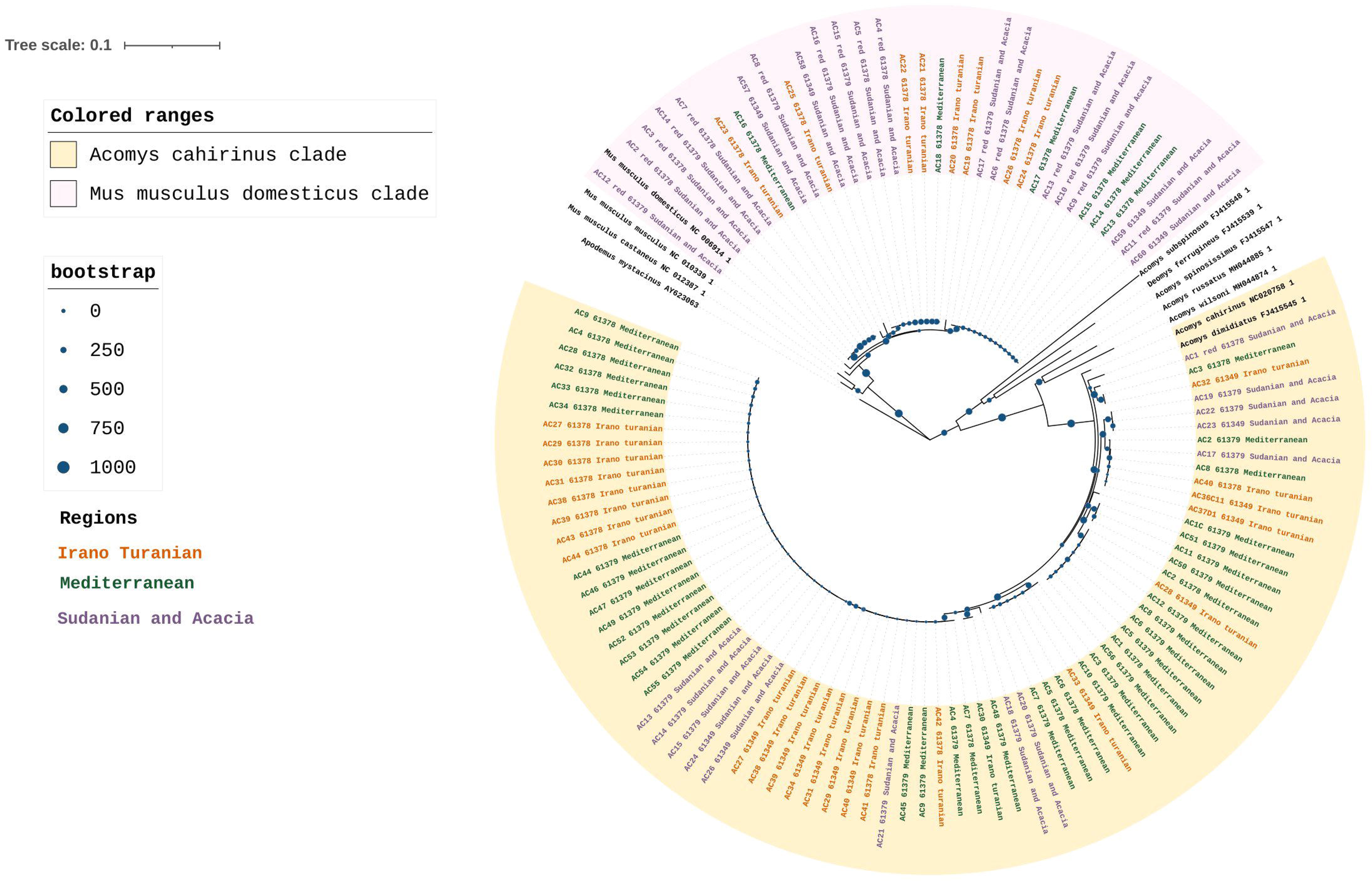
Rodent phylogeny. Maximum likelihood phylogenetic tree of the D-loop in *M. m. domesticus* and *A. cahirinus* s. l. Values for posterior likelihood and bootstrap support are shown for each node.

### Gut microbiome composition is associated with bioclimatic zone

16S rRNA gene amplicon sequencing of the rodent gut contents was performed to taxonomically characterize the gut microbiota composition. According to the species accumulation curve analysis, enough rodents were sampled to characterize the microbiome from the three climatic zones, with an ASV detection coverage of 84–96% based on Chao1 estimation (Figure 3A). A statistical analysis of the factors associated with microbiome composition using permutational multivariate analysis of variance (PerMANOVA) revealed that there were significant differences in the gut microbiota communities across the three bioclimatic zones (Table 2). PerMANOVA analysis showed that the microbiome profiles were associated bioclimatic zone (*p*= 0.001), which explained 17% of the total variation in the composition. Host species, sex, and weight were, however, not significantly associated with microbome composition (*p* > 0.05). In addition, there was no correlation between microbiome composition and host phylogeny as determined by Mantel test, either when all samples were considered together as well as when host-species specific comparisons were made (*A. cahirinus*: *p* = 0.18, R = 0.06; *M. m. domesticus*: *p* = 0.67, R = 0.03; *A. cahirinus* and *M. m. domesticus*: *p* = 0.002, R = 0.12).

**Figure 3.**
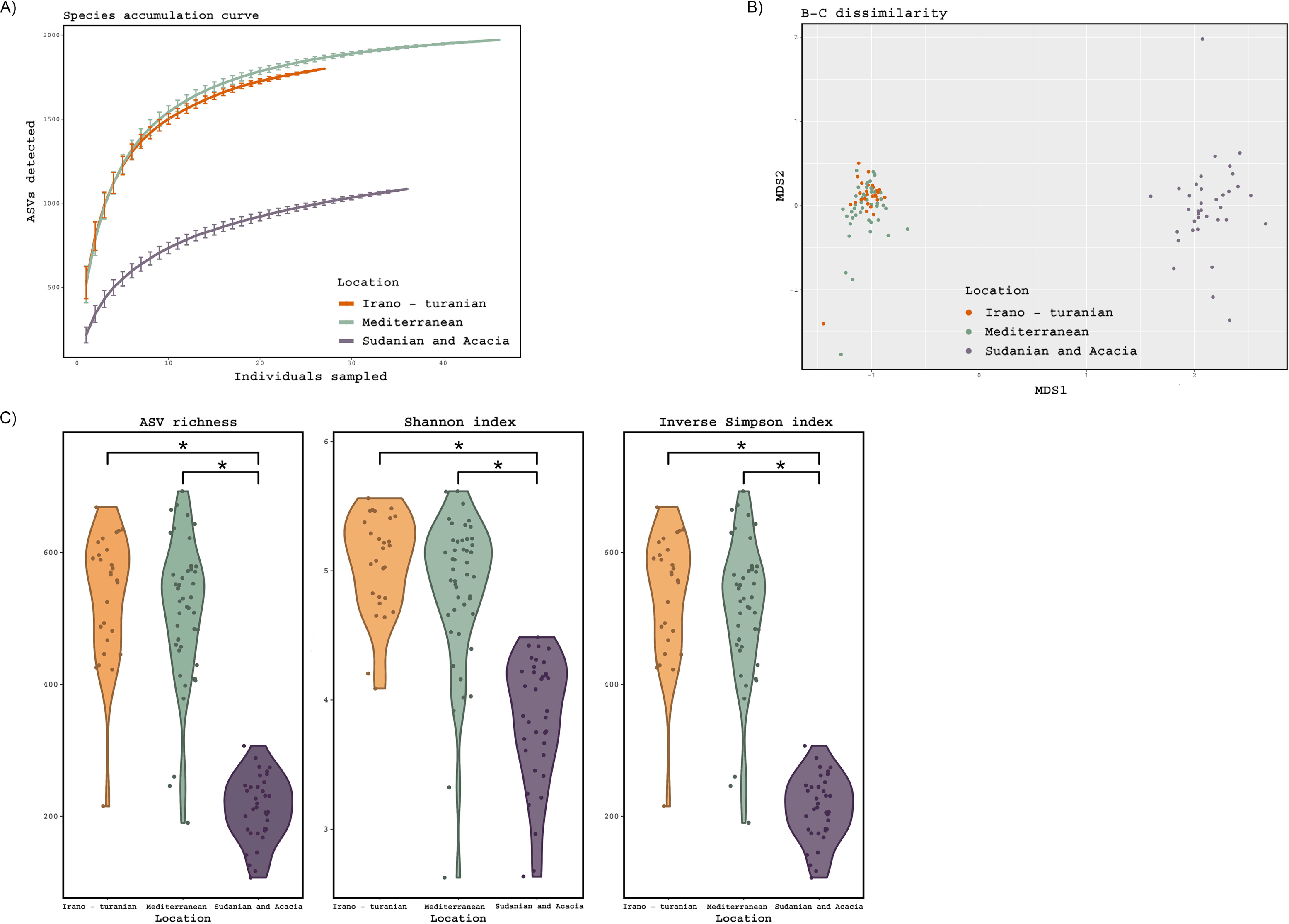
Diversity of the gut microbiome based on 16S rRNA gene amplicon analysis. (A) Species accumulation curve, determined at the level of ASVs. (B) Principal coordinate analysis ordinations based on Bray–Curtis distances of the gut microbiota of rodents collected from three bioclimatic zone, indicated by different colored dots. (C) Observed ASV richness, Shannon diversity, and inverse Simpson diversity across bioclimate zones. (ANOVA, p = 0.001; Tukey-HSD, p < 0.001). The threshold for significance is p = 0.05. The figure shows a black dot for the mean value and a colored region for the median value.

**Table 2.**
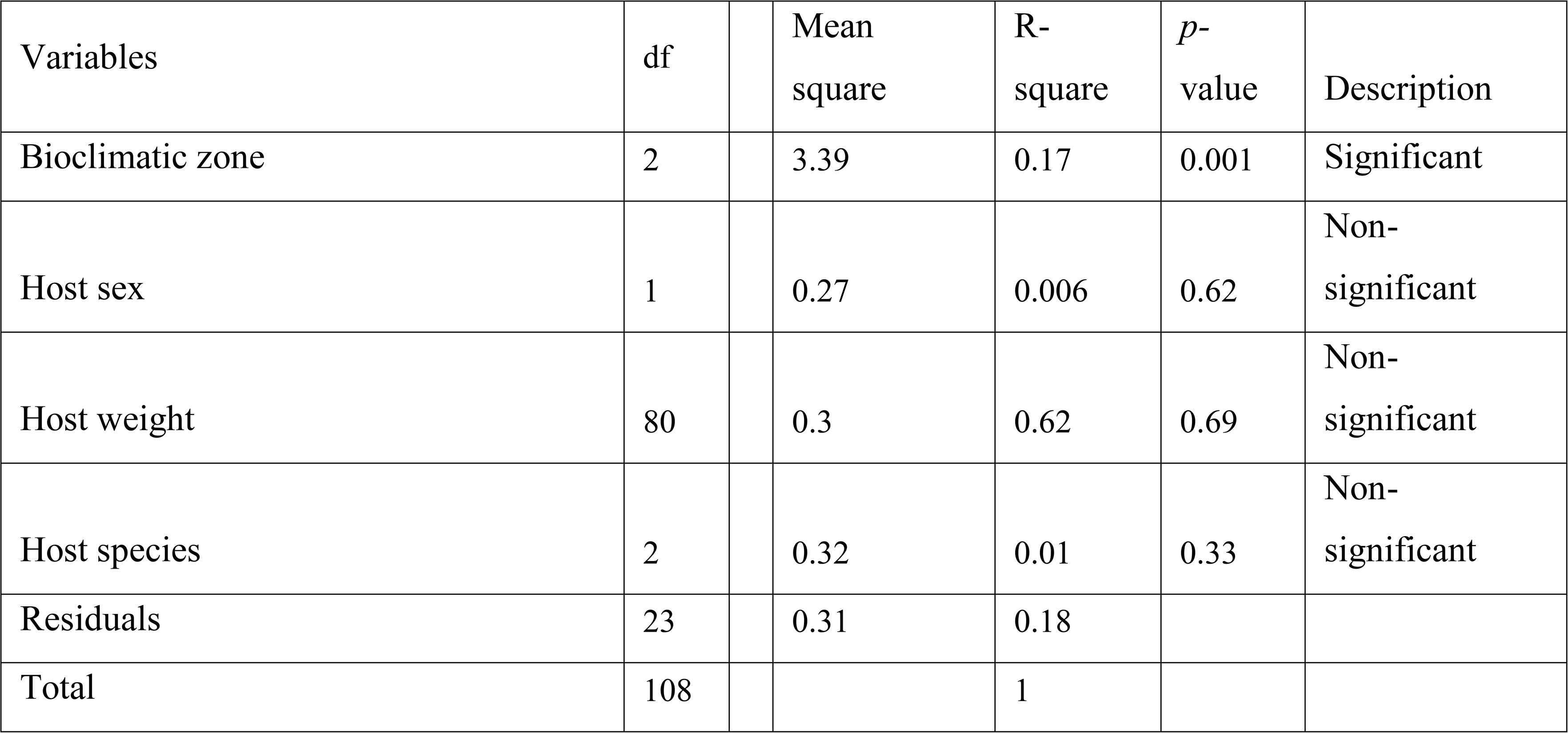
PerMANOVA analysis of the effect of host characteristics and the bioclimatic area on microbiota composition. Except for the bioclimatic zone type, none of the variables tested demonstrated any significant association.

Beta diversity ordination of the microbiome profiles also showed a distinct separation of samples from the Sudanian bioclimatic zone compared to the other two bioclimatic zones (Figure 3B). This was driven in part by a dramatically reduced richness and diversity of the gut microbiome from Sudanian bioclimatic zone rodents, as determined by observed ASV richness, Shannon diversity, and Inverse Simpson diversity indices (Figure 3C). However, the alpha diversity of rodents from the Mediterranean and Irano–Turanian bioclimatic zones were not significantly different.

### Taxanomic variation in the gut microbiota according to bioclimate zone

The gut microbiota of rodents from all bioclimatic zones was characterized by a high abundance of *Firmicutes* and *Bacteroidetes* phyla (Figure 4A), with unclassified *Lachnospiraceae*, *Lachnospiraceae* NK4A136 group, unclassified *Muribaculaceae*, and unclassified *Oscillospiraceae* being the predominant bacterial genera (Figure 4B). The phylum *Deferribacterota* was enriched in the Sudanian bioclimatic zone when compared to other bioclimatic zones (aldex.kw technique in ALDEx2 package, *p*≤0.05). At the genus level the samples from the Sudanian zone were also enriched in *Bacteroides* and *Oscillibacter* and depleted in *Alistipes* and unclassified *Oscillospiraceae* compared to samples from the Mediterranean and Irano-Turanian zones (*p* ≤0.05 for all taxa).

**Figure 4.**
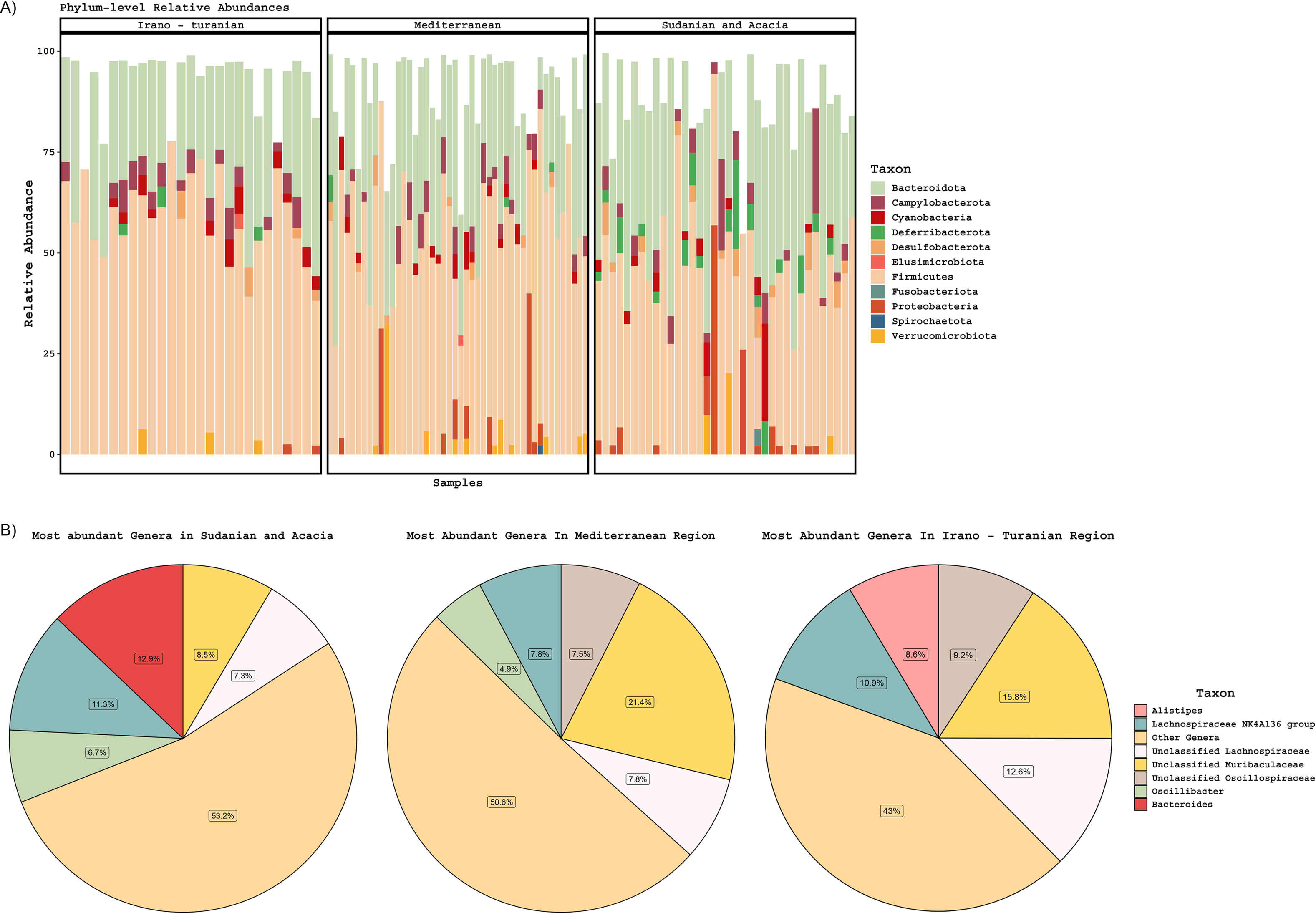
Taxonomic composition of gut microbiomes. (A) Stacked barplots of phylum-level diversity across the three climate zones. (B) The average abundance of the five most abundant genera is represented in a pie chart according to bioclimatic zone.

### Sudanian bioclimatic zone rodents have a unique microbiome gignature

Consistent with the alpha and beta diversity outcomes, differential ASV abundance analysis showed that the Mediterranean and Irano–Turanian bioclimatic zones had a greater number of differentially-abundant ASVs compared with the Sudanian bioclimatic zone than with each other (Figure 5A). Interestingly, some ASVs were more abundant in the lower diversity Sudanian samples, suggesting that Sudanian microbiomes are not just a subset of the diversity found in the other locations. To further evaluate the ASVs that were discriminative for the different climate zones, we used Random Forest machine learning to identify ASVs that were predictive of bioclimatic zones. Only common (> = 5 samples) and abundant ASVs (maximum relative abundance > = 1 %), which made up a total of 548 ASV, were included in the analysis. The algorithm identified 75 ASVs with a minimum and maximum frequency of 1% that occurred only in the Sudanian bioclimatic zone (Figure 5B, Supplementary Table 1). This analysis confirmed that the lower diversity Sudanian microbiomes do harbor unique diversity that is not merely a loss of diversity due to acclimatization to the Sudanian bioclimatic zone.

**Figure 5.**
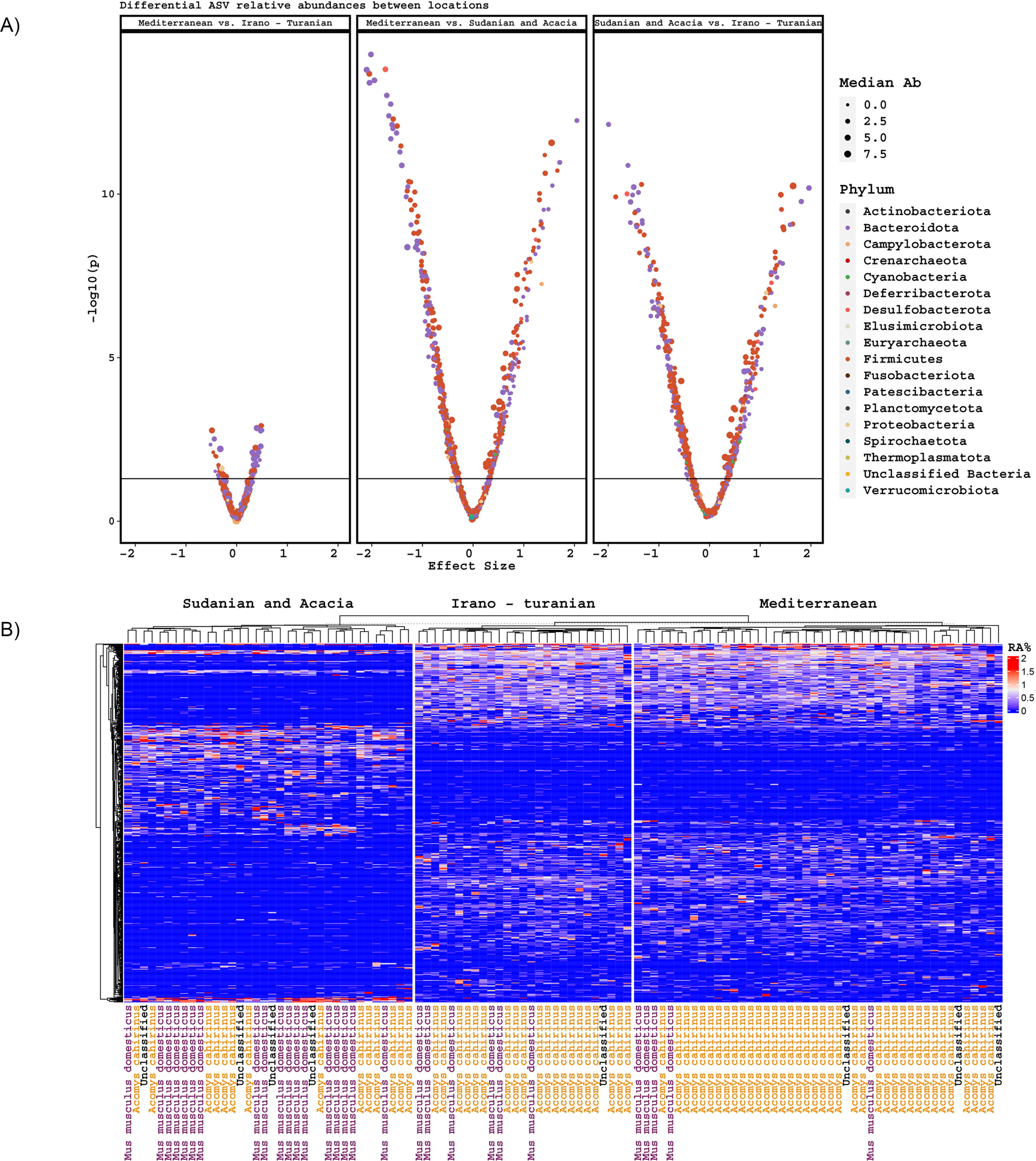
Differentially abundant ASVs. (A) Volcano plots of differentially abundant ASVs based on climatic zone, including effect size and p-value, and colored by phylum. (B) Heatmap of ASVs discriminative for climate zone, as determined by Random Forest machine learning.

### *18S* rRNA gene analysis reveals plant nutrition diversity

Analysis of 18S rRNA gene amplicon sequencing was hindered by the copious presence of rodent DNA. After removing these sequences, we were left with a relatively small number of reads putatively associated with either the intestinal microeukaryotes or dietary components. We used this data to perform a preliminary characterization of dietary plants. From the dataset, 26 *Embryophyceae* ASVs were identified, which are presumably part of rodent nutrition (Supplementary Table 2). We were unable to detect a difference in the richness or composition of *Embryophyceae* ASVs across bioclimatic zones despite trends for differences in average plant species composition (Figure 6, *p* = 0.2), though care must be taken in interpreting this as so few sequencing reads were recovered.

**Figure 6.**
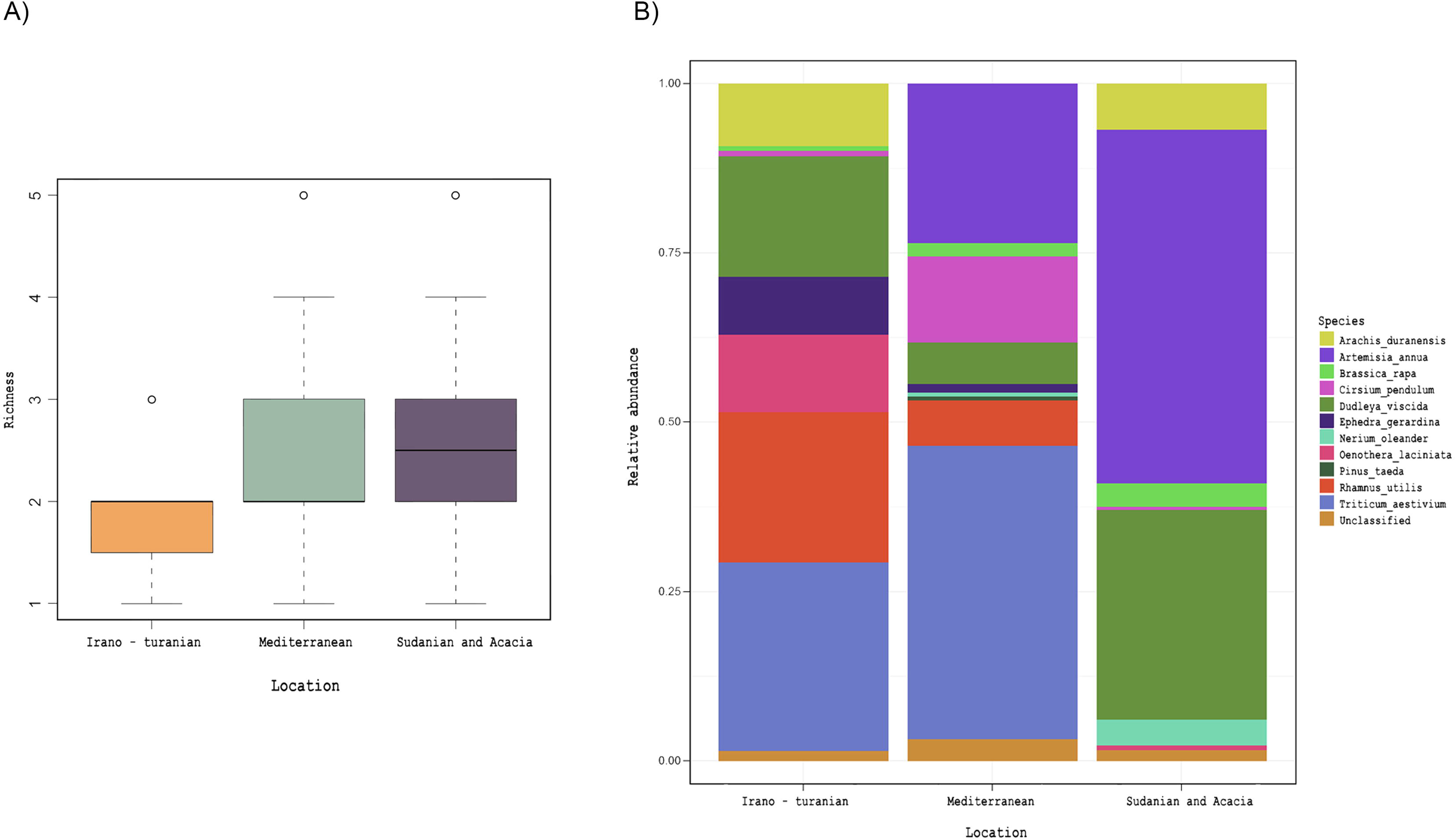
Plant diet, as inferred by 18S rRNA gene amplicon detection in rodent cecum. *Embryophyceae* ASVs obtained from 18S rRNA gene amplicon analysis. (A) Detected *Embryophyceae* richness across bioclimatic zones. (B) Average taxonomic composition of *Embryophyceae* across bioclimatic zones.

## Discussion

The evolution of phenotypic variation across phylogenies, including the interactions between species and their symbionts, is influenced by two factors, namely adaptive changes, such as responses to selective pressures, and phylogenetic stability, which is the similarity between species brought on by a shared evolutionary history. Therefore, it may be difficult to determine the relative contributions of these two elements to natural populations using comparative approaches. However, by examining tension zones, which are regions bordered by two different bioclimatic zones, we can determine how ecological and historical variables have changed the composition of the gut microbiome of conspecifics living in different environments. African GRVs have drawn the interest of scientists looking for untapped biodiversity and answers regarding the evolutionary past of the area. This study focused on the house mouse *M. m. domesticus* and spiny mouse *A. cahirinus*, which coexist in three distinct bioclimatic zones in the Jordanian portion of the GRV. Sequencing of the *D loop* region for phylogenetic analysis demonstrated that the two taxa under examination were clearly distinct from one another and that they had overlapping ranges. We studied the microbiota architecture of the two species but were unable to detect any significant interactions between host phylogeny and microbiota. The genetic differences that have emerged over the eight million years since these animals were separated are likely insufficient to distinguish their microbiomes beyond bioclimatic variances. These results have supported the findings of some earlier animal and rodent research, where it had not been possible to link the microbiota of the hosts to their evolutionary history (Teng et al., 2022; Griffiths et al., 2019; Baxter et al., 2015; Phillips et al., 2012; Ochman et al., 2010). Uncertainty exists over the minimum evolutionary time required to distinguish between the microbiota of two phylogenetically related species.

We hypothesized that animals collected from the same bioclimatic zone would be more likely to have a more similar microbiota than animals collected from different environments. Depending on the bioclimatic zone, we observed certain phenotypic variations between conspecifics, primarily a darker coat color and lower weight in the samples from the Sudanian zone. In this study, coat color variance brought on by melanin pigment synthesis in spiny and house mice was consistent with Gloger’s rule, which states that endothermic animals are darker in tropical climates than in other settings. Such phenotypic adaptability has already been noted in the house mouse, *Mus musculus* found throughout Asia (Lai et al., 2008), and in spiny mice on the tropical side of the GRV (Singaravelan et al., 2010). If we consider the microbiome as a source of phenotypic plasticity, this result suggests that climate plays a role as a selective factor that changes the structure and diversity of the microbiome. This will, in turn, influence the adaptive evolution of the host in bioclimatic zones.

The bioclimatic zones considered for this cohort occurred on an altitudinal gradient that varied in tandem with the ambient temperature gradient. Given that temperature and altitude are significant abiotic factors that shape the composition of animal populations and determine their adaptive arcs, the composition of the gut microbiota in *A. cahirinus* and *M. m. domesticus* is likely influenced by these factors. Bioclimatic areas can be classified as low altitude/hot (Sudanian), high altitude/cool to temperate (Mediterranean), or mid altitude/semi-temperate (Irano–Turanian). The Mediterranean and Irano–Turanian climatic zone with its moderate ambient temperature range/high to mid altitude was home to the greatest number of observed microbiome species based on the alpha diversity. Meanwhile, the Sudanian region, which has ambient temperatures that are typically 10 degrees higher and receive 10 times as much solar radiation, has the lowest number of observed microbiome species. The highest gut microbiota diversity was found in the Mediterranean and Irano–Turanian bioclimatic zones and the lowest was found in the Sudanian bioclimatic zone. Our differential abundance analysis has examined the impact of climate zones on microbiome makeup in various climatic regions. The differences in the abundance of each taxon between the Mediterranean and Irano– Turanian bioclimatic zones compared with the Sudanian site are noteworthy. To explain these findings, it is likely that variations in temperature and altitude shaped the makeup of the gut microbiomes of the rodent examined. In contrast to captive animals, wild rodents are constantly exposed to the combined thermal effects of their surroundings, including heat gain from sources such as solar radiation, long-wave radiation, conduction, and convection. High temperatures can increase beta diversity among host groups and disturb the stability of alpha diversity in the gut microbes of individual hosts (Sepulveda and Moeller, 2020b; Chevalier et al., 2015; Lan et al., 2017). The extreme temperatures in the Sudanian bioclimatic zone may be responsible for disrupting the stability of alpha diversity within the gut microbiomes of individual hosts and increasing beta diversity among the microbiomes within host populations.

Animals may find it challenging to release metabolic heat in a hot environment, such as the Sudanese climate zone, when there is a lower temperature gradient between the body core and the environment, which reduces the ability for body heat escape. In this study, similar temperature-induced microbiome responses were observed in heterospecific hosts across the Sudanese bioclimatic zone. This indicated that temperature-induced plasticity in the microbiome might be produced by conserved pathways. A similar notion has been mentioned in previous studies (Sepulveda and Moeller, 2020b). One way through which temperature-induced alterations in the gut microbiome can occur through the rapid response of animals to such circumstances, which is a reduction in feed intake (Berihulay et al., 2019). Given their higher basal metabolic rates per gram, small mammals, such as rodent, are thought to be more vulnerable to warmer conditions. In contrast to cold temperatures, these animals are beneficial. Various animal species, including mice, exhibit a steady decline in the relative abundance of Firmicutes with rising temperatures (Chevalier et al., 2015; Hylander and Repasky, 2019).

The data from our study are among the first to make characterize microbiome profiles with a view on temperature comparisons across various environments. However, for many animal taxa, we lack knowledge of the effects of these anticipated shifts on natural populations. High-temperature environments offer an opportunity to study the potential effects of ambient temperature on the composition and function of the gut microbiota with the potential to link this to global temperature regimes, which are expected to undergo rapid shifts in the next few decades.

According to previous data, disruption of the gut microbiome may be a mechanism by which altitude affects animal fitness in wild populations. This disruption is predominantly associated with an increase in microbiome diversity characterized by the presence of numerous obligate anaerobic bacteria induced by hypoxia at high elevations (Moreno-Indias et al., 2015; Zhang et al., 2018). Suzuki et al. (2019) identified a strong association between altitudinal changes and the alpha- and beta-diversity of the gut microbiome in house mice caused by taxon-specific variations (Suzuki et al., 2019). In line with these findings, greater alpha and beta diversities were found at higher altitude sites in this study in the Mediterranean and Irano–Turanian regions. However, the high-altitude populations studied here were from elevations between 1000 and 1500 m above sea level, where hypobaric hypoxia is unlikely to occur. Therefore, it is likely that rodent in these regions did not undergo significant selection because of hypoxic stress. The accompanying heterogeneity of gut microbiota can be explained by additional research on the physiological and genomic adaptability of animals living at high and low altitudes. In contrast, the Sudanian bioclimatic zone transect in our study included the lowest points on Earth, that is, the Dead Sea and Wadi Araba. Air has a slightly higher oxygen content in this zone because of its higher atmospheric pressure ranging from 3.3% in summer to 4.8% in winter. Lower microbial diversity and abundance in the Sudanian region may be related to variations in air oxygen levels linked to low altitudes. Although there is evidence to support short-term increases in oxygen abundance reducing microbial diversity in lab cultures (Shimizu and Benno, 2015; Comolli et al., 2023), it is still unclear if the makeup of the gut microbiota is reacting to the availability of atmospheric oxygen or whether it is a consequence of environmental host adaptation. Investigation of predictions regarding the impact of climatic zones using the Random Forest machine-learning technique showed that the Sudanian bioclimatic zone had distinct or novel microbial taxa that are not found among conspecifics inhabiting other bioclimatic zones. The specific functions of these bacteria may reflect significant variations in the abiotic parameters available. Diet and vegetation in the sampling zones may also be responsible for the observed differences in the microbial communities.

*A. cahirinus* and *M. m. domesticus* are omnivorous and eat snails, insects, seeds, and other plant matter (Amr et al., 2018a). Samples from all bioclimatic zones were abundant in *Firmicutes* and *Bacteroidetes* phyla, which is consistent with other studies of omnivores (Goertz et al., 2019; Nagpal et al., 2018). Conjointly, this outcome is in line with what is known about the mammalian gut containing a highly restricted collection of bacterial phyla that have adapted to the gastrointestinal tract environment (Thursby and Juge, 2017). Omnivores tend to have a greater proportion of *Bacteroidetes* to *Firmicutes* than other types of diets, such as lacto-ovo-vegetarians (Franco-de-Moraes et al., 2017), which is consistent with our findings. Although samples from all bioclimatic zones were abundant in *Firmicutes* and *Bacteroidetes*, the microbiome profiles from the Sudanian zone had a significantly higher abundance of *Deferribacterota*, which are prevalent in the healthy mouse gut (Chung et al., 2020), but also can expand during intestinal inflammation (Berry et al., 2015). The microbiome profiles from the Sudanian zone also showed significant changes in the microbiota composition of the genera of *Oscillibacter* and *Bacteroides,* which are known to metabolizes polysaccharides and oligosaccharides to deliver nutrients and vitamins to the host and other gut microbial inhabitants. There was a pronounced variation in the microbiome composition across the bioclimatic sites investigated (*p* ≤0.05). Our results are in line with studies on wild house mice, which discovered that geographic variation significantly affects the abundance of *Bacteroides* and *Lachnospiraceae* members (Goertz et al., 2019; Suzuki et al., 2019). Additionally, we observed a distinct divergence within the phylum *Firmicutes* between the studied bioclimatic zones, with *Alistipes* and unclassified *Oscillospiraceae* more abundant in Iranian and Mediterranean rodents and *Oscillibacter* in the Sudanian zone. Mice, ruminants, and humans frequently contain these *Firmicutes* taxa (Youssef et al., 2018; Yang et al., 2021), however, diet composition is considered to have an impact on their abundance (Youssef et al., 2018) (Pesoa et al., 2021). For instance, it has been found that mice fed a high-fat diet have an increased abundance of *Oscillibacter* in their feces (Lam et al., 2012). This raises the possibility that nutrition influences *Oscillibacter* fluctuation between the bioclimatic zones. Our research suggests that the sustained nutritional and environmental changes on rodents cause shifts in their intestinal microbiome. This could result from coevolutionary selective pressure acting on both the host and the microbiome.

There were no differences in the abundance of bacterial genera between the Irano-Turanian and Mediterranean bioclimatic zones, although some differentially abundant ASVs were detected. The topographic map of the study site (Figure 1) depicts the Irano–Turanian region as a narrow strip encircling the Mediterranean region. Rodents could move easily between the two sites because the Irano–Turanian region was occasionally considered a segment of the Mediterranean region. Therefore, it is likely that the overlap in the ranges of bioclimatic zones caused individual mice to migrate between the two sites, resulting in a population that was mixed or admixed and had a comparable microbiome signature. This finding may be our second piece of evidence in favor of the theory that the phylosymbiosis pattern is because of environmental determinants rather than genetic diversity and familial ties. Local patterns of genetic diversity in our phylogenetic analysis did not correlate with geography, and the heterospecific and genetically distinct *A. cahirinus* and *M. m. domesticus*, which live in the same locations, have similar microbiomes. However, we cannot rule out the involvement of host phylogeny. The extent to which the environment and the host’s genetic heritage or phylogeny influence the pattern of phylosymbiosis has long been controversial. More detailed genetic and environmental data are required to differentiate these two alternatives. Settings such as the contrasting abiotic parameters being studied and the genetic distance between the studied hosts, may affect how patterns of phylosymbiosis are interpreted, particularly when animal hosts are evaluated. Host species account for less variance in the gut microbiota of rats in comparison to biogeography factors (Linnenbrink et al., 2013; Teng et al., 2022; Baxter et al., 2015) with host species, suggesting commonalities in gut microbiota among species (Doms et al., 2022; Nagpal et al., 2018; Weinstein et al., 2021). However, the host environment and genetics interact to shape phylosymbiosis (Wang et al., 2022; Griffiths et al., 2019). The core bacteria identified in this study may have important effects on health and fitness. However, some rodent microbiomes may contain zoonotic agents. *Campylobacter*, for example, can cause numerous human and animal diseases, including gastroenteritis (Song et al., 2021). Additionally, the high abundance of *Proteobacteria* in some samples may be related to dysbiosis in hosts with inflammatory or metabolic illnesses (Rizzatti et al., 2017).

Whether dietary changes are a driver of the composition of the microbiota, while controlling for other variables such as climate, is one of the prominent gaps in the earlier phylosymbiosis studies, with relatively few exceptions. According to Baxter et al. (2015), diet was unrelated to the intra- and inter-species heterogeneity in mouse microbiota (Baxter et al., 2015). Quantifying the impact of nutrition alone on wild microbiomes is challenging. We characterized the dietary content of each *A. cahirinus* and *M. m. domesticus* sample using 18S rRNA metabarcoding analysis to provide estimate of the relative richness and diversity of the components in the rodent diets. We focused on plant composition (*Embryophyceae*). This analysis demonstrated that the bioclimatic zone was not significantly correlated with the number of plants and seeds. However, the composition of plants in the gut was correlated. Based on this, convergent diets may encourage the same microbiota in heterospecific hosts that shared the same niche, such as *A. cahirinus* and *M. m. domesticus*. In laboratory studies, nutritional manipulation has created selective pressures on the microbiome that are stronger than those caused by host genetics (Huda et al., 2022). The changes in the microbiota observed in this study may have been influenced by diet. This will aid our understanding of phylosymbiosis patterns in natural populations.

## Conclusion

In this study, rodent gut microbiota composition was associated the abiotic factors in the bioclimatic zones. This suggests that neutral assembly and dispersal did not cause the microbiota composition of the heterospecific host species to diverge. In contrast, the conspecific hosts in this study included microbiomes that varied depending on the bioclimatic zone from which they were mainly assembled. Host phylogeny was not an important determinant of the microbiota composition. The host species *A. cahirinus* and *M. m. domesticus* have been shown to be able to thrive in various environmental conditions, which reflects their capacity to change their diet. Concomitant changes in the gut microbiota may play a role in improved metabolic adaptation and improved energy extraction. However, future work is needed to uncover the functional implications of the phylosymbiosis.

## Conflict of Interest

*The authors declare that the research was conducted in the absence of any commercial or financial relationships that could be construed as a potential conflict of interest*.

## Author Contributions

**E. Al-khlifeh:** Provided concepts and ideas, structured the research, defined the intellectual content, searched for relevant literature, conducted experimental studies, collected samples, and wrote the manuscript.

**S. Khadem:** Produced Fig.s data acquisition, data analysis, and manuscript writing.

**D. Berry:** Data acquisition, data analysis, and manuscript revision.

**B. Hausmann:** Experimental studies.

All authors have reviewed and approved the manuscript.

## Funding

Funding provided by Al-Balqa Applied University-Deanship of scientific research (DSR), grant Id **DSR-2020#226**

## Acknowledgments

We acknowledge Jordan’s Royal Society of Nature Conservation (RSNC), for allowing access to sampling site and providing a map of three adjacent bioclimatic zones from the DANA biosphere on south of Jordan (Figure1).

## Data Availability Statement

16S and 18S rRNA gene amplicon MiSeq sequence data has been deposited in the NCBI Short Read Archive under the BioProject accession PRJNA992969. Rodent and mitochondrial *D-loop* sequence data has been deposited in GenBank under the BioProject ID PRJNA992969.

## Notes

### Competing Interest Statement

The authors have declared no competing interest.

